# Hypothalamic connectivities and self-evaluated aggression in young adults

**DOI:** 10.1101/2024.09.26.615292

**Authors:** Yuxing Jared Yao, Yu Chen, Chiang-Shan R. Li

## Abstract

**Introduction:** The hypothalamus plays a pivotal role in supporting motivated behavior, including aggression. Previous work suggested differential roles of the medial hypothalamus (MH) and lateral hypothalamus (LH) in aggressive behaviors, but little is known about how their resting-state functional connectivity (rsFC) may relate to aggression in humans.

**Methods:** We employed the data from the Human Connectome Project (HCP) and examined the rsFC’s of LH and MH in 745 young adults (393 women). We also explored sex differences in the rsFC’s. We processed the imaging data with published routines and evaluated the results of voxel-wise regression on aggression score, as obtained from Achenbach Adult Self Report, with a corrected threshold.

**Results:** The analysis revealed significant rsFC between the LH and clusters in the middle temporal and occipital gyri across all subjects and in the thalamus for men, both in negative correlation with aggression score. Slope test confirmed sex differences in the correlation between LH-thalamus rsFC and aggression score. No significant rsFC was observed for MH.

**Conclusions:** These findings suggest a role of LH rsFCs and sex differences in LH-thalamus rsFC in the manifestation of aggression in humans. The findings highlight the need for further research into sex-specific neural pathways in aggression and other related behavioral traits of importance to mental health.

## 1. Introduction

### 1.1 The hypothalamus and motivated behavior

Connected with a wide swath of brain regions (Rizzi et al., 2021), the hypothalamus plays a vital role in regulating many physiological functions, including feeding, mating, and sleep, as well as a wide range of motivated behavior, including aggression (Mignot et al., 2002; Neary et al., 2004; Williams et al., 2001). For instance, studies have demonstrated hypothalamic projections to the amygdala and medial prefrontal cortex (mPFC), referred to as the ventral system, in the relay of processed sensory information to support motivated behavior (Risold et al., 1997; Valenstein et al., 1970). Medial prefrontal cortical connections to the lateral hypothalamus (LH) are essential for learning of cued feeding (Allen & Cechetto, 1993; Petrovich et al., 2005), with stimulation in mPFC eliciting LH activity and feeding in sated rats (Mena et al., 2013). In humans, imaging studies highlighted the roles of the hypothalamus in regulating socioemotional responses (Caria, 2023) and of the hypothalamic amygdala and ventromedial PFC (vmPFC) connectivity in supporting decision making (Averbeck & Murray, 2020).

### 1.2 Hypothalamic circuits and aggression

The hypothalamus is implicated in animal studies in the perception of threat cues, representation of an aggressive state, and execution of aggressive behaviors (Hashikawa et al., 2017). Previous work has also distinguished the roles of the LH and medial hypothalamus (MH) in aggression. The LH appears to play an outsized role in generating predatory (Smith et al., 1970) and intraspecific (Tulogdi et al., 2015) aggression whereas the MH is more active in defensive aggression in response to external stimulation. For instance, in mice, electrical stimulation of LH induced intraspecific attacks with the presence of a subordinate but not a dominant male (Koolhaas, 1978). In mice, stimulation of mPFC glutamatergic projection terminals in the LH elicited violent bites toward conspecifics (Roeling et al., 1994). However, another study provided evidence discounting the roles of the LH by showing that optogenetic stimulation of the MH but not LH induced attack behaviors in male mice (Lin et al., 2011).

Falkner et al. described a MH-midbrain pathway organizing social signals and tensing jaw muscles during aggressions in mice (Falkner et al., 2020). Other studies implicated the MH-thalamus and spinal cord circuit in reactive aggression (Gouveia et al., 2019; Haller, 2018). Amygdala’s projection to the hypothalamus has also been studied in relation to aggressive behaviors (Canteras et al., 1995). Optogenetic stimulation showed that synaptic potentiation in the amygdala-MH pathway contributed to aggression in rats in response to traumatic stress (Nordman et al., 2020). Studies in rhesus monkeys revealed inhibitory PFC projections to the hypothalamus (de Boer et al., 2015) and its role in regulating behavioral responses evoked externally (Rempel-Clower & Barbas, 1998). Together, the evidence suggests a role of MH in the generation of aggression, particularly defensive aggression in response to environmental stimulation. In view of these contrasting findings and an earlier work characterizing distinct rsFC of the LH and MH (Kullmann et al., 2014), it would seem important to investigate the roles of LH and MH circuit function and dysfunction in motivated behavior.

The hypothalamus has also long been implicated in the generation and regulation of aggressive behaviors in humans, with bilateral lesions of the LH documented as a treatment for outbursts of aggression in the mid-20th century (Rizzi et al., 2021). More recently, deep brain stimulation in the treatment of patients with intermittent explosive disorder also included the hypothalamus as a target (Rizzi et al., 2021). A positron emission tomography imaging study reported significant lower metabolism in the right hypothalamus of men engaged in domestic violence, relative to control participants (George et al., 2004). A functional magnetic resonance imaging (fMRI) study associated activation of the limbic and subcortical areas, including the hypothalamus, in reaction to physical provocation (Fanning et al., 2017). Hypothalamic activations are also found during threats of pain and conspecific attacks (Bertram et al., 2022), social threats, and witnessing of aggression towards strangers (Caria & Dall, 2022). These studies support the roles of the hypothalamus in aggressive responses; however, little is known of hypothalamic circuit function and dysfunction in manifesting individual differences and whether the roles of MH and LH are distinguishable in aggression.

### 1.3 Sex differences in aggression

Empirical evidence suggests sex differences in the manifestation of aggression, with males being more aggressive than females, particularly with respect to physical aggression (Bjorkqvist, 2018; Tieger, 1980). Boys exhibited more physical aggressive behaviors while girls showed more indirect aggression, including social exclusion and gossiping (Rosvall et al., 2012). The sex differences may be rooted in genetics and associated with circulating testosterone. An fMRI study reported that, when provoked, men tended to have higher left amygdala activations than women. Men were also found to demonstrate a positive association between orbitofrontal cortex (OFC), rectal gyrus, and anterior cingulate cortex (ACC) activity and their tendency to respond aggressively, while women displayed a negative association during a modified Taylor Aggression Task (Repple et al., 2018).

Animal work supports sex differences in aggressive behaviors mediated by hypothalamus. A study revealed modulatory effects of hormone expression in the MH and its circuits on neural development in the prenatal period and subsequent aggressive behaviors in male but not female mice (Yang et al., 2024). Arginine-vasopressin (AVP) is a neuropeptide crucial in the regulation of aggressive behavior. The injection of AVP into the hypothalamus stimulated aggression among male but reduced aggression in female hamsters (Gutzler et al., 2010). Similar sex-dependent effects were found in humans, with intranasal administration of AVP enhancing affiliative responses in women in face of social stimuli while enhancing agonistic responses in men (Thompson et al., 2006). These findings suggest important sex differences in the neural mechanisms of aggression (Terranova et al., 2017).

### 1.4 The present study

We investigated how hypothalamic rsFC’s may vary with individual aggression, using data from the Human Connectome Project (HCP). Seed-based rsFC’s reflect functional organization of the neural circuits (Fair et al., 2007; Zhang et al., 2017) and capture individual variation in behavioral traits and psychopathology (Li et al., 2014; Woodward & Cascio, 2015; Zhang et al., 2017). Previous studies have characterized rsFC of the habenula in people with intermittent explosive disorder (Gan et al., 2019) and cortical rsFCs in children and adolescents with disruptive behaviors (Werhahn et al., 2023). Given the literature of potentially differentiable roles of the LH and MH and sex differences in aggression, we investigated the rsFC’s of the LH and MH as well as in men and women separately.

## 2. Methods

### 2.1 Dataset

We have obtained permission from the HCP to use both open and restricted access data. We employed data from the 1200 Subjects Release (S1200), which includes behavioral and 3T MRI from 1206 young adults (1113 with structural MR scans). A group of 220 subjects were excluded due to excessive head movement (translation >2 mm or rotation >2° in any dimension) and/or problematic image quality after normalization.

Participants completed the Achenbach Adult Self Report (ASR) (Achenbach & Rescorla, 2003) for the assessment of aggressive behaviors. We used ASR raw scores in our analyses. The ASR includes 8 syndromal scales (Anxious/Depressed; Withdrawn; Somatic Complaints; Thought Problems; Attention Problems; Aggressive Behaviors; Rule-breaking Behaviors; and Intrusive Problems) and 6 DSM-oriented scales (Depressive; Anxiety; Somatic; Avoidant Personality; Attention Deficit/Hyperactivity; and Antisocial Personality problems), with each scored: 0-Not True, 1-Somewhat or Sometimes True, 2-Very or Often True. In addition to these item scores, ASR provided 8 additional raw scores: Inattention and Hyperactivity/Impulsivity subscales, Other Problems, Critical Items, Internalizing Problems (Anxious/Depressed Syndromes, Withdrawn Syndromes, and Somatic Complaints), Externalizing Problems (Aggressive Behaviors, Rule-breaking Behaviors, and Intrusive Problems), TAO sum (Thought Problems, Attention Problems, and Other Problems), and Total Problems (all problems). Thus, there were a total of 22 ASR measures, with higher scores indicating more symptoms. We focused on the raw score of aggressive behaviors in the current study. Eight participants were excluded from the analyses due to missing ASR scores.

Participants were also assessed with the Semi-Structured Assessment for the Genetics of Alcoholism (SSAGA), an instrument designed to evaluate physical, psychological, and social manifestations of alcoholism and related disorders (Bucholz et al., 1994). As in our previous study (Li et al., 2022), we performed a principal component analysis on the 15 interrelated drinking metrics and identified one principal component (PC1) with an eigenvalue > 1 that accounted for 49.75% of the variance. Alcohol use may contribute to aggression (Chen et al., 2024; Li et al., 2022; Li et al., 2021); thus, to control for the effect of alcohol use, we included drinking PC1 as a covariate in all analyses. Data from 38 participants were excluded because of missing alcohol use scores. The final sample comprised a total of 745 subjects (393 females; age: 28.47±3.71 years).

The key demographic and clinical measures are shown in Error! Reference source not found.. Women were significantly older than men (t=-6.718, p<0.001). Men showed higher ASR raw aggression scores (t=2.367, p=0.018) and higher drinking severity PC1 (t=9.869, p<0.001) than women. Chi-square test showed no significant difference in race composition between sexes (χ^2^=2.718, p=0.743).

**Table 1:**
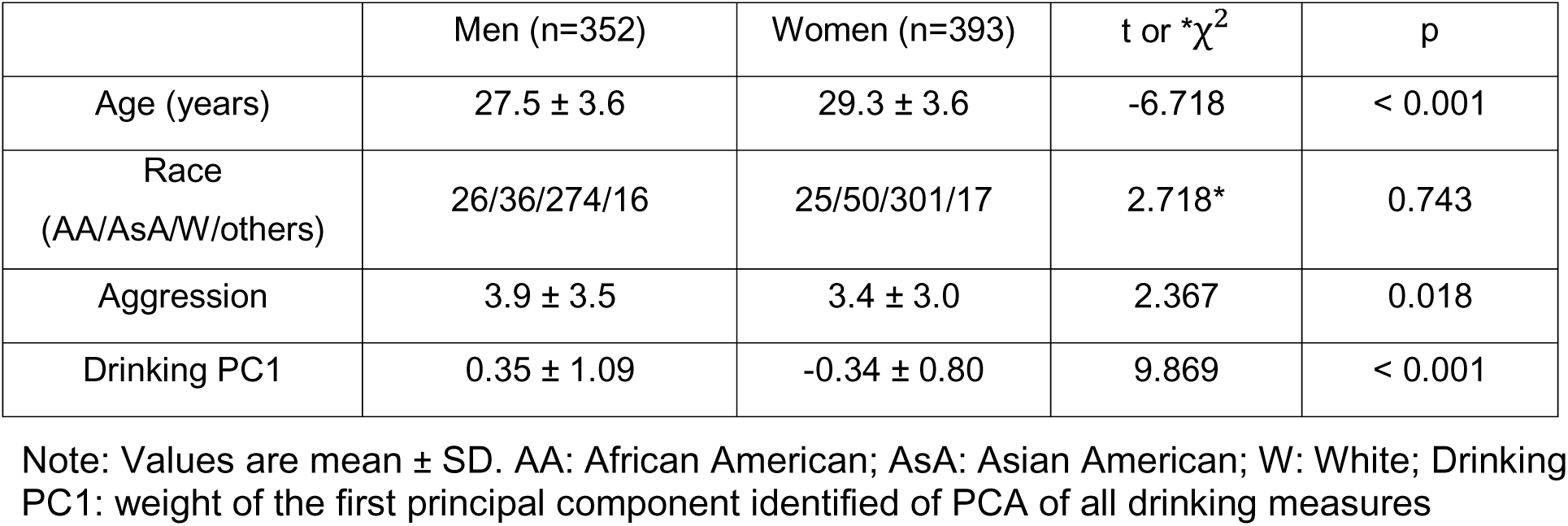
Demographic and clinical measures in men and women.

| | Men (n=352) | Women (n=393) | t or $\chi^2$ | p |
| --- | --- | --- | --- | --- |
| Age (years) | 27.5 ± 3.6 | 29.3 ± 3.6 | -6.718 | < 0.001 |
| Race<br>(AA/AsA/W/others) | 26/36/274/16 | 25/50/301/17 | 2.718* | 0.743 |
| Aggression | 3.9 ± 3.5 | 3.4 ± 3.0 | 2.367 | 0.018 |
| Drinking PC1 | 0.35 ± 1.09 | -0.34 ± 0.80 | 9.869 | < 0.001 |

Note: Values are mean ± SD. AA: African American; AsA: Asian American; W: White; Drinking PC1: weight of the first principal component identified of PCA of all drinking measures

### 2.2 MRI data acquisition and imaging data processing

All imaging data were acquired on a customized Siemens 3T Skyra with a standard 32-channel Siemens receiver head coil and a body transmission coil. T1-weighted high-resolution structural images were acquired using a 3D MPRAGE sequence with 0.7 mm isotropic resolution (FOV = 224 mm, matrix = 320, 256 sagittal slices, TR = 2400 ms, TE = 2.14 ms, TI = 1000 ms, FA = 8°) and used to register resting state fMRI (rsfMRI) data to a standard brain space. The rsfMRI data were collected in two sessions, using gradient-echo echo-planar imaging (EPI) with 2.0 mm isotropic resolution (FOV = 208×180 mm, matrix = 104×90, 72 slices, TR = 720 ms, TE = 33.1 ms, FA = 52°, multi-band factor = 8). Physiological data (i.e., cardiac, and respiratory signals) associated with each fMRI scan were also acquired, using a standard Siemens pulse oximeter placed on a digit and a respiratory belt placed on the abdomen. These physiological signals were sampled equally at 400 Hz (∼288 samples per frame). More details of the data collection procedures can be found in the HCP S1200 Release Reference Manual. Within each session, oblique axial acquisitions alternated between phase encoding in a right-to-left (RL) direction in one run and phase encoding in a left-to-right (LR) direction in the other run. Each run lasted 14.4 min (1200 frames), with associated physiological signals sampled at 400 Hz (∼288 samples/frame).

We preprocessed the resting-state fMRI data with SPM8 as described earlier (Chen et al., 2022; Chen & Li, 2023), including realignment, coregistration, normalization, smoothing, regression of confounding signals (i.e., white matter, cerebrospinal fluid, whole-brain mean, and physiological signals), temporal band-pass filtering, and scrubbing. In the current study, only the first session (two runs: LR and RL) of resting-state fMRI data were used and processed with SPM8. Images of each participant were first realigned (motion corrected), and a mean functional image volume was constructed from the realigned image volumes. These mean images were co-registered with the high-resolution structural MPRAGE image and then segmented for normalization with affine registration followed by nonlinear transformation. The normalization parameters determined for the structural volume were then applied to the corresponding functional image volumes for each participant. Afterwards, the images were smoothed with a Gaussian kernel of 4 mm at Full Width at Half Maximum. White matter and cerebrospinal fluid signals, whole-brain mean signal, and physiological signals were regressed out to reduce spurious BOLD variances and to eliminate cardiac- and respiratory-related artifacts. A temporal band-pass filter (0.009 Hz < *f* < 0.08 Hz) was also applied to the time course to obtain low-frequency fluctuations, as in our prior work (Zhang & Li, 2018).

Lastly, to further eliminate global motion-related artifacts, a “scrubbing” method was applied. Specifically, frame-wise displacement given by *FD*(*t*) = |Δ*d*_x_(*t*)| + |Δ*d*_y_(*t*)| + |Δ*d*_z_(*t*)| + |Δα(*t*)| + |Δβ(*t*)| + |Δγ(*t*)| was computed for every time point *t*, where (*d*_x_, *d*_y_, *d*_z_) and (α, β, γ) are the translational and rotational movements, respectively (Power et al., 2012). Moreover, the root mean square variance of the differences (DVARS) in % BOLD intensity *I*(*t*) between consecutive time points across brain voxels, was computed as: *DVARS*(*t*) = sqrt(|*I*(*t*) – *I*(*t*-1)|^2^), where the brackets indicate the mean across brain voxels. Following previous HCP studies (Chen et al., 2021; Li et al., 2019), we marked volumes with FD > 0.2 mm or DVARS > 75 as well as one frame before and two frames after these volumes as outliers (censored frames). Uncensored segments of data lasting fewer than five contiguous volumes were also labeled as censored (Li et al., 2019). A total of 6 smokers who had both BOLD runs with more than half of the frames scrubbed were removed from further analyses.

### 2.3 Seed-based correlation for MH and LH rsFC

The seed masks of MH (two spheres of 2 mm in radius, centered at x = ±4, y = −2, z = −12) and LH (two spheres of 2 mm in radius, centered at x = ±6, y = −9, z = −10) were obtained from the WFU Pick Atlas (http://fmri.wfubmc.edu/software/pickatlas) (Breen et al., 2016) as in our previous studies (Zhang et al., 2018). We computed the whole-brain rsFC and estimated the correlation coefficient *r* between the average time course of all voxels of the seed and the time courses of all other voxels of the brain for individual participants. To assess and compare the rsFC, we converted these image maps, which were not normally distributed, to z score maps by Fisher’s z transform (Jenkins & Watts, 1968): 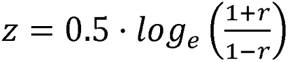. The Z maps were used in group random effect analyses.

### 2.4 Group data analyses

We first performed a one-sample t test of whole-brain rsFCs of the LH and MH and paired sample t test of LH and MH rsFC, so we could compare the results with those reported previously (Kullmann et al., 2014; Zhang et al., 2018). To address our specific aims, we performed whole-brain regression against trait aggression on the rsFCs of LH and MH in all subjects, with age, sex, race, and drinking PC1 as covariates, and in male and female subjects separately, with age, race, and drinking PC1 as covariates. We evaluated the results at voxel p < 0.005, uncorrected, in combination with a cluster p < 0.05, corrected for family-wise error (FWE), on the basis of Gaussian random field theory, as implemented in SPM, following current reporting standards (Poldrack et al., 2008). At this threshold, one-sample *t* tests yielded large, contiguous clusters, and thus we also evaluated one-sample *t* tests at voxel *p* < 0.05 FWE-corrected. We also conducted paired t-test at voxel *p* < 0.05 FWE-corrected to compare the rsFCs between LH and MH. In addition to reporting the peak voxel *Z* value, we computed the effect size by approximating Cohen’s *d* from the *t*-statistics, *t* value and degrees of freedom (*df*), using the expression 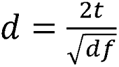 (Howell, 2013).

For the clusters revealed in the whole-brain analyses in men or women alone, we used the masks of the ROIs to compute for all subjects the β estimates of their rsFCs with LH and MH. We then employ slope tests to examine sex differences in the correlations between the hypothalamic rsFCs and aggression. Note that this analysis did not represent “double-dipping.” The clusters were identified with a statistical threshold; a cluster identified in men may have just missed the threshold in women, and vice versa. Thus, slope tests were needed to confirm sex differences.

## 3. Results

### 3.1 Whole-brain hypothalamic rsFC

We conducted a one-sample t test of whole-brain rsFC of the LH and MH for all subjects. Both LH and MH showed extensive connectivities encompassing the frontal, parietal, temporal, and occipital cortices, insula, cerebellum, and subcortical regions (Error! Reference source not found.). We also conducted paired t test of whole-brain rsFCs of the LH vs. MH. The result showed that LH had significantly higher rsFC than MH in the thalamus, middle and inferior frontal gyri, insular cortex, cerebellum, supramarginal cortex, inferior parietal gyrus, dorsal anterior cingulate cortex (ACC), supplementary motor cortex, putamen, and caudate, while MH had significantly higher rsFC than LH in medial and superior frontal gyri, medial orbitofrontal cortex, hippocampus, parahippocampal area, superior temporal gyrus, middle and superior occipital gyri, precuneus, angular gyrus and cingulate gyrus. These findings were consistent with those of Kullman and colleagues (Kullmann et al., 2014).

**Figure 1:**
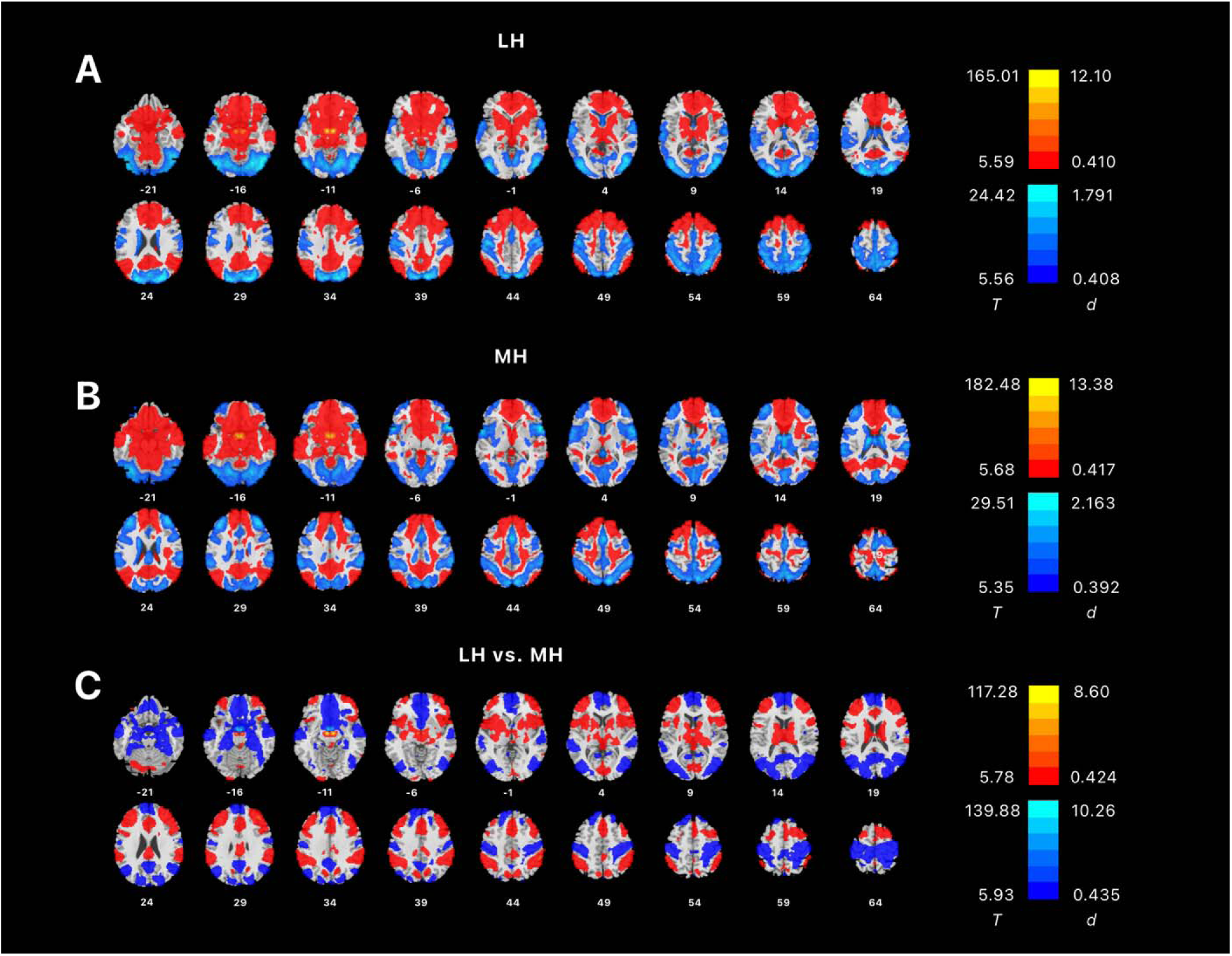
Whole-brain resting-state connectivity of (A) LH and (B) MH shown in one-sample t-tests, evaluated at voxel p < 0.05 corrected for family-wise error (FWE). Warm and cool color each shows positive and negative rsFC. (C) showed paired t-test of LH vs. MH rsFCs, evaluated at voxel p < 0.05, FWE corrected. Color bars show voxel T value and Cohen’s d. Warm/cool color: in (A) and (B) positive/negative resting state functional connectivity, and in (C), positive/negative differences in resting state functional connectivity between LH and MH.

### 3.2 LH and MH rsFC correlates of aggression: whole-brain analysis

Across all subjects, whole-brain regression of LH rsFCs on aggression score showed negative rsFC with the left middle temporal gyri (x=-64, y=-14, z=-14, 1552 voxels, Z=3.78) and right middle occipital gyri (x=28, y=-80, z=-18, 2968 voxels, Z=4.29). Regression of MH rsFC evaluated at the same threshold did not reveal significant clusters. In male subjects alone, we found negative rsFC of LH with a cluster in the right thalamus (x=9, y=-19, z=17, 1768 voxels, Z=3.75) but no significant clusters for MH. No significant clusters were identified for women for either LH or MH rsFCs in correlation with aggression score. These results are shown in **Figure 2**.

**Figure 2.**
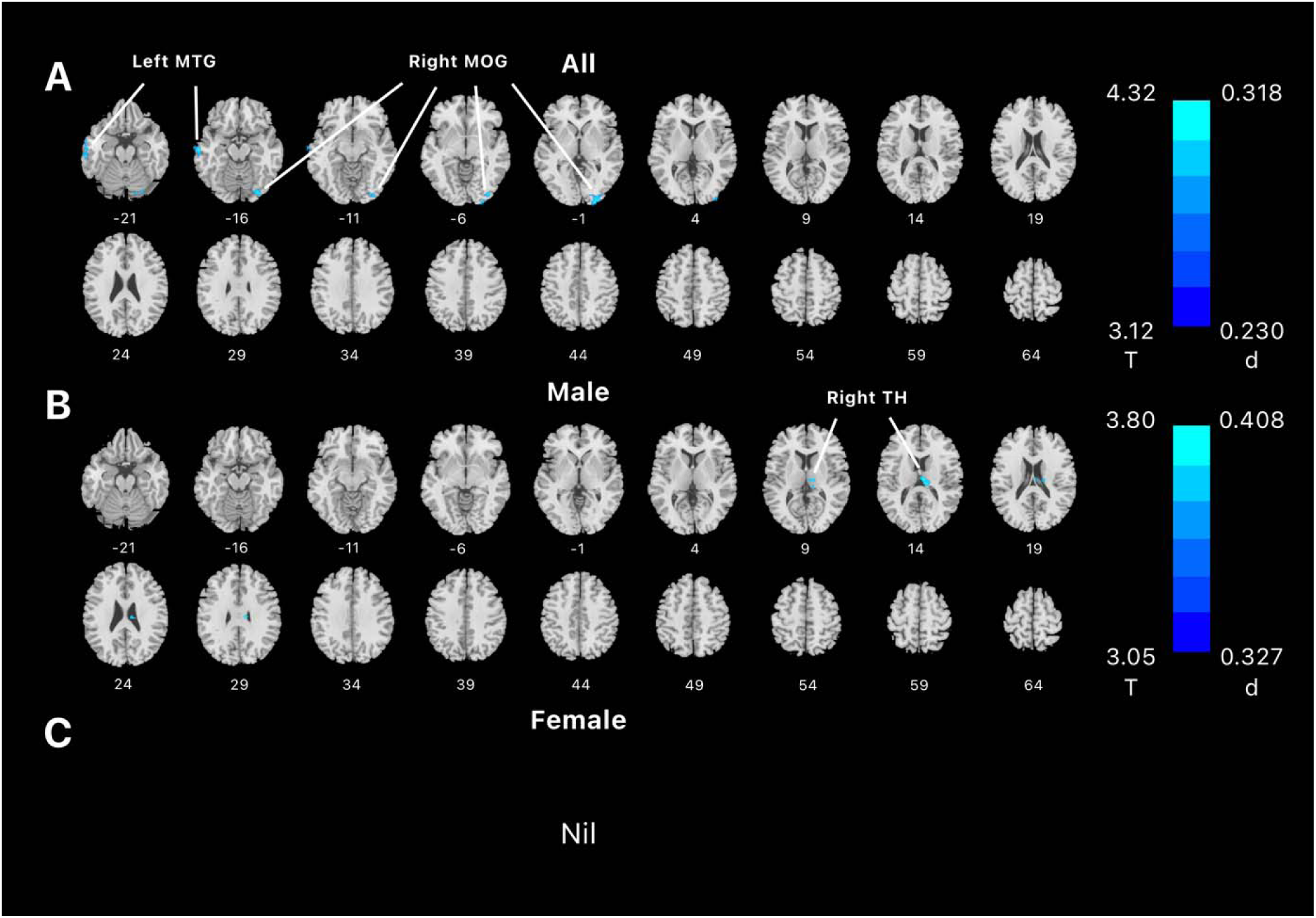
Regression of whole-brain resting-state functional connectivity of lateral hypothalamus on aggression score, evaluated at voxel p < 0.005 uncorrected and cluster p < 0.05 FWE corrected in (**A**) all, (**B**) male, and (**C**) female subjects. Color bar shows voxel T value and Cohen’s d. Cool color reflected negative correlation. Nil: no significant findings; MTG: middle temporal gyrus; MOG: Middle occipital gyrus; TH: thalamus.

Because the thalamus cluster was identified in men alone, we performed slope test to examine sex differences in the correlation of LH-thalamus rsFC and aggression. This was needed because the cluster was identified with a threshold, and the thalamus cluster may have just missed the threshold in women. The correlation between LH-thalamus rsFC and aggression scores was significant, as expected (r = 0.0697, p < 0.0001). However, the correlation between LH-thalamus rsFC and aggression scores was not significant in women (r = 0.0006, p = 0.6206). Slope test confirmed the sex difference (t = 3.2388, p = 0.0013). **Figure 3** shows the scatter plots and regression lines for men and women separately.

**Figure 3:**
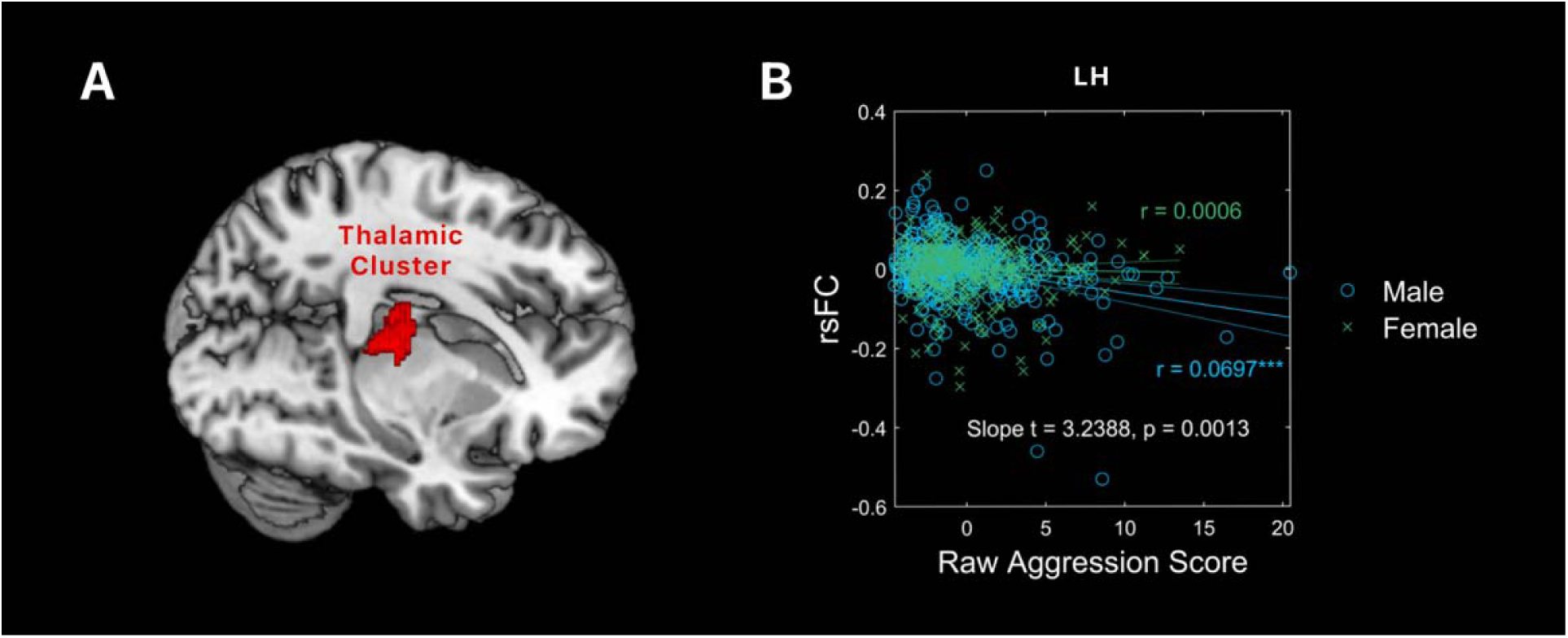
(A) The thalamic cluster isolated from whole-brain regression of lateral hypothalamus rsFC on aggression score in men. (B) A scatter plot showing the correlations of rsFC between LH and the thalamic cluster with aggression in men and women, with age, race, and drinking PC1 as covariates. Residuals are presented here. Each data point represents one subject. Solid lines represent the regressions: blue for male and green for female. ***p < 0.0001.

## 4. Discussion

The current study examined how resting-state functional connectivity (rsFC) of medial hypothalamus (MH) and lateral hypothalamus (LH) varies with individual aggression. Whole-brain analyses revealed LH rsFC’s with the right occipital cortex and left temporal cortex across all subjects and with the thalamus in men alone, both in negative correlation with aggression. We discussed the main findings below.

### 4.1 LH and MH rsFC correlates of aggression

Aggressive behaviors are defined as those carried out with the intention to cause harm to others (Krahé, 2020). The HCP assessed aggression with the ASR, which captures primarily intraspecific aggression but not reactive aggression as provoked by external stimuli. As discussed earlier, animal studies support a role of the LH and MH in intraspecific and reactive aggression, respectively. Thus, our findings of LH but not MH connectivities are broadly consistent with this literature. Whether or how MH rsFCs may be involved in aggression would likely need to be investigated in laboratory paradigms, e.g., the Ultimatum game (Harsanyi, 1961), that involve interaction with others.

Across all subjects, LH-temporal and occipital cortex rsFC’s were significantly correlated with aggression score. Both temporal and occipital cortices are implicated in a meta-analysis of fMRI studies of individuals with a history of aggression, who, relative to controls, showed higher activation in the temporal and occipital cortices, along with the hippocampus, parahippocampal gyrus, and amygdala, during aggression-eliciting tasks (Nikolic et al., 2022). An earlier review associated temporal lobe dysfunction with unwarranted, poorly directed anger outbursts, with left hemispheric lateralization among aggression-prune subjects (Potegal, 2012). The author proposed a temporo-frontal model of aggression, where visual and auditory threat perceptions are processed in the temporal cortex, which projects to the vmPFC and OFC for higher-level evaluation and eventual motivated behavior (Potegal, 2012).

### 4.2 Hypothalamus thalamus connectivity and aggression

A cluster in right thalamus showed significant negative rsFC with LH in association with higher aggression in males but not in females, with the sex difference confirmed by a slope test. The cluster comprised the mediodorsal nucleus of thalamus (MD) (Behrens et al., 2003). In animal studies, the MD and paraventricular nucleus of thalamus (PVT) were frequently discussed together with respect to their roles in aggression (Mai & Majtanik, 2018). Early animal studies showed that stimulation in the thalamus could both induce and suppress aggressive behaviors in mice (Bandler, 1971). Furthermore, suppressive stimulation in the thalamus blocked the facilitation of aggression induced by hypothalamic stimulation, whereas hypothalamic stimulation had no effect on thalamus stimulation-induced aggression (Bandler, 1971). This result indicated that aggression elicited by hypothalamus stimulation is contingent on the thalamic activity, in accord with a potential modulatory role of the thalamus in the generation of aggressive behaviors. One study using calcium imaging-based fiber photometry showed LH GABAergic innervations of PVT neurons, providing a neurochemical pathway to support hypothalamic influences of thalamic functions (Otis et al., 2019; Vertes et al., 2022). The latter study can perhaps be considered together with the current findings of hypothalamic thalamic rsFC in negative correlation with aggression.

Earlier animal studies have implicated the PVT and MD in aggression. For instance, in cats, electrical stimulation of the thalamus elicited attacks similar to the quiet biting attack elicited by stimulation of the LH (Bandler & Flynn, 1974). Another posited that the thalamus integrated motor and sensory information and modulated aggressive behavior, with clinical reports of patients with thalamic lesions manifesting pain insensibility and uncontrolled aggression (Andy et al., 1975). One study reported that rats bearing MD lesions were significantly less aggressive (Ferreira et al., 1987). An fMRI study reported activations in the LH, basal ganglia, as well as various thalamic areas, including the PVT and medial thalamus, in male rats in the company of their female cage mate presented with a novel male intruder (Ferris et al., 2008). A more recent model of aggression also included the thalamus, along with the amygdala and PFC, as a key component in the bottom-up circuit modulating aggression in mice (Hashikawa et al., 2017).

### 4.3 Sex difference in aggression

The results show a significant sex difference in hypothalamic rsFC with the thalamus in link with individual aggression. Men vs. women showed a significantly more negative rsFC between the hypothalamus and thalamus in correlation with aggression scores. This finding suggests hypothalamic-thalamic connectivity as a neural marker specifically of male aggression. An earlier study showed that, contrary to increases in aggressive behaviors in male Syrian hamsters following injection of arginine-vasopressin (AVP) in the hypothalamus, female hamsters exhibited reduction in aggression (Gutzler et al., 2010). The results suggest that the same hormones, such as AVP, engage different neural pathways in males and females, leading to distinct behaviors in aggression. The relation between hypothalamus and the thalamus, PVT and/or MD specifically, among males can be further studied.

### 4.4 Limitations and conclusion

Several limitations of the study need to be considered. First, the HCP comprised largely a neurotypical sample. Therefore, the current findings should be considered as specific to this population. More studies are needed to investigate whether the findings may hold in people with mental conditions that manifest aggression. Second, as described earlier, the hypothalamus is involved in a variety of motivated behavior. Here, we related the connectivity findings to subjectively rated aggression. It remains to be seen how the hypothalamic circuits are involved in laboratory-based “real-life” aggressive behavior, an issue that may be particularly important in light of the roles of the MH in aggression. Likewise, studies should be conducted to address the roles of hypothalamus connectivity in other motivated behaviors, including reward and punishment processing, in link with aggression. Finally, aggression manifests differently between males and females. Behavioral paradigms that are “sex relevant” may be considered to further investigate the hypothalamic correlates of sex differences in aggression.

In summary, we demonstrated hypothalamic connectivity correlates of aggression and sex differences in the LH circuits in aggression. The current findings complement the imaging literature of aggression in humans and may help in the development of targeted interventions of aggressive behaviors.

## CRediT authorship contribution statement

Yuxing Jared Yao: Software, Data Curation, Formal Analysis, Writing – Original Draft, Writing – Review & Editing, Visualization. Yu Chen: Conceptualization, Methodology, Writing – Original Draft, Writing – Review & Editing. Chiang-Shan R. Li: Conceptualization, Methodology, Writing – Original Draft, Writing – Review & Editing, Supervision, Funding Acquisition.

## Acknowledgements

This study is supported by NIH grant DA051922. The NIH is otherwise not responsible for the conceptualization of the study, data collection and analysis, or in the decision in publishing the results.

## Data availability, Ethics declarations, and Consent to participate

We have obtained permission from the Human Connectome Project (HCP) to use the Open and Restricted Access data for the current study. Data were provided by the WU-Minn Consortium (Principal Investigators: David Van Essen and Kamil Ugurbil; 1U54MH091657) funded by the 16 NIH Institutes and Centers that support the NIH Blueprint for Neuroscience Research; and by the McDonnell Center for Systems Neuroscience at Washington University. The HCP young-adult data is publicly available at https://www.humanconnectome.org/study/hcp-young-adult/.

## References

Achenbach, T. M., & Rescorla, L. A. (2003). Manual for the ASEBA adult forms & profiles. In: Burlington, VT: University of Vermont.

Allen, G. V., & Cechetto, D. F. (1993). Functional and Anatomical Organization of Cardiovascular Pressor and Depressor Sites in the Lateral Hypothalamic Area. 2. Ascending Projections. Journal of Comparative Neurology, 330(3), 421–438. 10.1002/cne.903300310

Andy, O. J., Giurintano, L., Giurintano, S., & McDonald, T. (1975). Thalamic modulation of aggression. Pavlov J Biol Sci, 10(2), 85–101. 10.1007/BF03001153

Averbeck, B. B., & Murray, E. A. (2020). Hypothalamic Interactions with Large-Scale Neural Circuits Underlying Reinforcement Learning and Motivated Behavior. Trends Neurosci, 43(9), 681–694. 10.1016/j.tins.2020.06.006

Bandler, R. J. (1971). Direct chemical stimulation of the thalamus: Effects on aggressive behavior in the rat. Brain research., 26(1), 81–93. 10.1016/0006-8993(71)90544-0

Bandler, R. J., Jr., & Flynn, J. P. (1974). Neural pathways from thalamus associated with regulation of aggressive behavior. Science, 183(4120), 96–99. 10.1126/science.183.4120.96

Behrens, T. E., Johansen-Berg, H., Woolrich, M. W., Smith, S. M., Wheeler-Kingshott, C. A., Boulby, P. A., Barker, G. J., Sillery, E. L., Sheehan, K., Ciccarelli, O., Thompson, A. J., Brady, J. M., & Matthews, P. M. (2003). Non-invasive mapping of connections between human thalamus and cortex using diffusion imaging. Nat Neurosci, 6(7), 750–757. 10.1038/nn1075

Bertram, T., Hoffmann Ayala, D., Huber, M., Brandl, F., Starke, G., Sorg, C., & Mulej Bratec, S. (2022). Human threat circuits: Threats of pain, aggressive conspecific, and predator elicit distinct BOLD activations in the amygdala and hypothalamus. Front Psychiatry, 13, 1063238. 10.3389/fpsyt.2022.1063238

Bjorkqvist, K. (2018). Gender differences in aggression. Curr Opin Psychol, 19, 39–42. 10.1016/j.copsyc.2017.03.030

Breen, D. P., Nombela, C., Vuono, R., Jones, P. S., Fisher, K., Burn, D. J., Brooks, D. J., Reddy, A. B., Rowe, J. B., & Barker, R. A. (2016). Hypothalamic volume loss is associated with reduced melatonin output in Parkinson’s disease. Movement Disorders, 31(7), 1062–1066.

Canteras, N. S., Simerly, R. B., & Swanson, L. W. (1995). Organization of projections from the medial nucleus of the amygdala: a PHAL study in the rat. Journal of Comparative Neurology, 360(2), 213–245. 10.1002/cne.903600203

Caria, A. (2023). A Hypothalamic Perspective of Human Socioemotional Behavior. Neuroscientist, 10738584221149647. 10.1177/10738584221149647

Caria, A., & Dall, O. G. (2022). Functional Neuroimaging of Human Hypothalamus in Socioemotional Behavior: A Systematic Review. Brain Sci, 12(6). 10.3390/brainsci12060707

Chen, Y., Chaudhary, S., Li, G., Fucito, L. M., Bi, J., & Li, C. R. (2024). Deficient sleep, altered hypothalamic functional connectivity, depression and anxiety in cigarette smokers. Neuroimage Rep, 4(1). 10.1016/j.ynirp.2024.100200

Chen, Y., Dhingra, I., Chaudhary, S., Fucito, L., & Li, C. R. (2022). Overnight Abstinence Is Associated With Smaller Secondary Somatosensory Cortical Volumes and Higher Somatosensory-Motor Cortical Functional Connectivity in Cigarette Smokers. Nicotine Tob Res, 24(12), 1889–1897. 10.1093/ntr/ntac168

Chen, Y., & Li, C. R. (2023). Overnight Abstinence, Ventrostriatal-Insular Connectivity, and Tridimensional Personality Traits in Cigarette Smokers. Journal of Integrative Neuroscience, 22(3), 1–8.

Chen, Y., Li, G., Ide, J. S., Luo, X., & Li, C. R. (2021). Sex differences in attention deficit hyperactivity symptom severity and functional connectivity of the dorsal striatum in young adults. Neuroimage: Reports, 1(2), 100025. 10.1016/j.ynirp.2021.100025

de Boer, S. F., Olivier, B., Veening, J., & Koolhaas, J. M. (2015). The neurobiology of offensive aggression: Revealing a modular view. Physiol Behav, 146, 111–127. 10.1016/j.physbeh.2015.04.040

Fair, D. A., Schlaggar, B. L., Cohen, A. L., Miezin, F. M., Dosenbach, N. U. F., Wenger, K. K., Fox, M. D., Snyder, A. Z., Raichle, M. E., & Petersen, S. E. (2007). A method for using blocked and event-related fMRI data to study “resting state” functional connectivity. Neuroimage, 35(1), 396–405. 10.1016/j.neuroimage.2006.11.051

Falkner, A. L., Wei, D., Song, A., Watsek, L. W., Chen, I., Chen, P., Feng, J. E., & Lin, D. (2020). Hierarchical Representations of Aggression in a Hypothalamic-Midbrain Circuit. Neuron, 106(4), 637–648 e636. 10.1016/j.neuron.2020.02.014

Fanning, J. R., Keedy, S., Berman, M. E., Lee, R., & Coccaro, E. F. (2017). Neural Correlates of Aggressive Behavior in Real Time: a Review of fMRI Studies of Laboratory Reactive Aggression. Curr Behav Neurosci Rep, 4(2), 138–150. 10.1007/s40473-017-0115-8

Ferreira, A., Dahlof, L. G., & Hansen, S. (1987). Olfactory mechanisms in the control of maternal aggression, appetite, and fearfulness: effects of lesions to olfactory receptors, mediodorsal thalamic nucleus, and insular prefrontal cortex. Behav Neurosci, 101(5), 709–717, 746. 10.1037//0735-7044.101.5.709

Ferris, C. F., Stolberg, T., Kulkarni, P., Murugavel, M., Blanchard, R., Blanchard, D. C., Febo, M., Brevard, M., & Simon, N. G. (2008). Imaging the neural circuitry and chemical control of aggressive motivation. BMC Neurosci, 9, 111. 10.1186/1471-2202-9-111

Gan, G., Zilverstand, A., Parvaz, M. A., Preston-Campbell, R. N., Uquillas, F. D., Moeller, S. J., Tomasi, D., Goldstein, R. Z., & Alia-Klein, N. (2019). Habenula-prefrontal resting-state connectivity in reactive aggressive men - A pilot study. Neuropharmacology, 156. ARTN107396 10.1016/j.neuropharm.2018.10.025

George, D. T., Rawlings, R. R., Williams, W. A., Phillips, M. J., Fong, G., Kerich, M., Momenan, R., Umhau, J. C., & Hommer, D. (2004). A select group of perpetrators of domestic violence: evidence of decreased metabolism in the right hypothalamus and reduced relationships between cortical/subcortical brain structures in position emission tomography. Psychiatry Res, 130(1), 11–25. 10.1016/S0925-4927(03)00105-7

Gouveia, F. V., Hamani, C., Fonoff, E. T., Brentani, H., Alho, E. J. L., de Morais, R., de Souza, A. L., Rigonatti, S. P., & Martinez, R. C. R. (2019). Amygdala and Hypothalamus: Historical Overview With Focus on Aggression. Neurosurgery, 85(1), 11–30. 10.1093/neuros/nyy635

Gutzler, S. J., Karom, M., Erwin, W. D., & Albers, H. E. (2010). Arginine-vasopressin and the regulation of aggression in female Syrian hamsters (Mesocricetus auratus). Eur J Neurosci, 31(9), 1655–1663. 10.1111/j.1460-9568.2010.07190.x

Haller, J. (2018). The Role of the Lateral Hypothalamus in Violent Intraspecific Aggression-The Glucocorticoid Deficit Hypothesis. Front Syst Neurosci, 12, 26. 10.3389/fnsys.2018.00026

Harsanyi, J. C. (1961). On the rationality postulates underlying the theory of cooperative games. Journal of Conflict Resolution, 5, 179–196. 10.1177/002200276100500205

Hashikawa, Y., Hashikawa, K., Falkner, A. L., & Lin, D. (2017). Ventromedial Hypothalamus and the Generation of Aggression. Front Syst Neurosci, 11, 94. 10.3389/fnsys.2017.00094

Howell, D. C. (2013). Statistical methods for psychology (Eigth edition. ed.). Wadsworth. https://www.vlebooks.com/vleweb/product/openreader?id=ARuskin&accId=9137456&isbn=9781133713272

Jenkins, G. M., & Watts, D. G. (1968). Spectral analysis and its applications. Holden-Day.

Koolhaas, J. M. (1978). Hypothalamically induced intraspecific aggressive behaviour in the rat. Exp Brain Res, 32(3), 365–375. 10.1007/BF00238708

Krahé, B. (2020). The social psychology of aggression. Routledge.

Kullmann, S., Heni, M., Linder, K., Zipfel, S., Haring, H. U., Veit, R., Fritsche, A., & Preissl, H. (2014). Resting-state functional connectivity of the human hypothalamus. Hum Brain Mapp, 35(12), 6088–6096. 10.1002/hbm.22607

Li, C. S. R., Ide, J. S., Zhang, S., Hu, S., Chao, H. H., & Zaborszky, L. (2014). Resting state functional connectivity of the basal nucleus of Meynert in humans: In comparison to the ventral striatum and the effects of age. Neuroimage, 97, 321–332. 10.1016/j.neuroimage.2014.04.019

Li, G., Chen, Y., Chaudhary, S., Tang, X., & Li, C. R. (2022). Loss and Frontal Striatal Reactivities Characterize Alcohol Use Severity and Rule-Breaking Behavior in Young Adult Drinkers. Biol Psychiatry Cogn Neurosci Neuroimaging, 7(10), 1007–1016. 10.1016/j.bpsc.2022.06.001

Li, G., Chen, Y., Tang, X., & Li, C. R. (2021). Alcohol use severity and the neural correlates of the effects of sleep disturbance on sustained visual attention. J Psychiatr Res, 142, 302–311. 10.1016/j.jpsychires.2021.08.018

Li, J., Kong, R., Liégeois, R., Orban, C., Tan, Y., Sun, N., Holmes, A. J., Sabuncu, M. R., Ge, T., & Yeo, B. T. T. (2019). Global signal regression strengthens association between resting-state functional connectivity and behavior. Neuroimage, 196, 126–141. 10.1016/j.neuroimage.2019.04.016

Lin, D., Boyle, M. P., Dollar, P., Lee, H., Lein, E. S., Perona, P., & Anderson, D. J. (2011). Functional identification of an aggression locus in the mouse hypothalamus. Nature, 470(7333), 221–226. 10.1038/nature09736

Mai, J. K., & Majtanik, M. (2018). Toward a Common Terminology for the Thalamus. Front Neuroanat, 12, 114. 10.3389/fnana.2018.00114

Mena, J. D., Selleck, R. A., & Baldo, B. A. (2013). Mu-Opioid Stimulation in Rat Prefrontal Cortex Engages Hypothalamic Orexin/Hypocretin-Containing Neurons, and Reveals Dissociable Roles of Nucleus Accumbens and Hypothalamus in Cortically Driven Feeding. Journal of Neuroscience, 33(47), 18540–18552. 10.1523/Jneurosci.3323-12.2013

Mignot, E., Taheri, S., & Nishino, S. (2002). Sleeping with the hypothalamus: emerging therapeutic targets for sleep disorders. Nat Neurosci, 5 Suppl, 1071-1075. 10.1038/nn944

Neary, N. M., Goldstone, A. P., & Bloom, S. R. (2004). Appetite regulation: from the gut to the hypothalamus. Clin Endocrinol (Oxf), 60(2), 153–160. 10.1046/j.1365-2265.2003.01839.x

Nikolic, M., Pezzoli, P., Jaworska, N., & Seto, M. C. (2022). Brain responses in aggression-prone individuals: A systematic review and meta-analysis of functional magnetic resonance imaging (fMRI) studies of anger- and aggression-eliciting tasks. Prog Neuropsychopharmacol Biol Psychiatry, 119, 110596. 10.1016/j.pnpbp.2022.110596

Nordman, J. C., Ma, X., Gu, Q., Potegal, M., Li, H., Kravitz, A. V., & Li, Z. (2020). Potentiation of Divergent Medial Amygdala Pathways Drives Experience-Dependent Aggression Escalation. J Neurosci, 40(25), 4858–4880. 10.1523/JNEUROSCI.0370-20.2020

Otis, J. M., Zhu, M., Namboodiri, V. M. K., Cook, C. A., Kosyk, O., Matan, A. M., Ying, R., Hashikawa, Y., Hashikawa, K., Trujillo-Pisanty, I., Guo, J., Ung, R. L., Rodriguez-Romaguera, J., Anton, E. S., & Stuber, G. D. (2019). Paraventricular Thalamus Projection Neurons Integrate Cortical and Hypothalamic Signals for Cue-Reward Processing. Neuron, 103(3), 423–431 e424. 10.1016/j.neuron.2019.05.018

Petrovich, G. D., Holland, P. C., & Gallagher, M. (2005). Amygdalar and prefrontal pathways to the lateral hypothalamus are activated by a learned cue that stimulates eating. Journal of Neuroscience, 25(36), 8295–8302. 10.1523/Jneurosci.2480-05.2005

Poldrack, R. A., Fletcher, P. C., Henson, R. N., Worsley, K. J., Brett, M., & Nichols, T. E. (2008). Guidelines for reporting an fMRI study. Neuroimage, 40(2), 409–414.

Potegal, M. (2012). Temporal and frontal lobe initiation and regulation of the top-down escalation of anger and aggression. Behav Brain Res, 231(2), 386–395. 10.1016/j.bbr.2011.10.049

Power, J. D., Barnes, K. A., Snyder, A. Z., Schlaggar, B. L., & Petersen, S. E. (2012). Spurious but systematic correlations in functional connectivity MRI networks arise from subject motion. Neuroimage, 59(3), 2142–2154. 10.1016/j.neuroimage.2011.10.018

Rempel-Clower, N. L., & Barbas, H. (1998). Topographic organization of connections between the hypothalamus and prefrontal cortex in the rhesus monkey. Journal of Comparative Neurology, 398(3), 393–419. 10.1002/(sici)1096-9861(19980831)398:3<393::aid-cne7>3.0.co;2-v

Repple, J., Habel, U., Wagels, L., Pawliczek, C. M., Schneider, F., & Kohn, N. (2018). Sex differences in the neural correlates of aggression. Brain Struct Funct, 223(9), 4115–4124. 10.1007/s00429-018-1739-5

Risold, P. Y., Thompson, R. H., & Swanson, L. W. (1997). The structural organization of connections between hypothalamus and cerebral cortex. Brain Res Brain Res Rev, 24(2-3), 197–254. 10.1016/s0165-0173(97)00007-6

Rizzi, M., Gambini, O., & Marras, C. E. (2021). Posterior hypothalamus as a target in the treatment of aggression: From lesioning to deep brain stimulation. Handb Clin Neurol, 182, 95–106. 10.1016/B978-0-12-819973-2.00007-1

Roeling, T. A. P., Veening, J. G., Kruk, M. R., Peters, J. P. W., Vermelis, M. E. J., & Nieuwenhuys, R. (1994). Efferent Connections of the Hypothalamic Aggression Area in the Rat. Neuroscience, 59(4), 1001–1024. 10.1016/0306-4522(94)90302-6

Rosvall, K. A., Bergeon Burns, C. M., Barske, J., Goodson, J. L., Schlinger, B. A., Sengelaub, D. R., & Ketterson, E. D. (2012). Neural sensitivity to sex steroids predicts individual differences in aggression: implications for behavioural evolution. Proc Biol Sci, 279(1742), 3547–3555. 10.1098/rspb.2012.0442

Smith, D. E., King, M. B., & Hoebel, B. G. (1970). Lateral hypothalamic control of killing: evidence for a cholinoceptive mechanism. Science, 167(3919), 900–901. 10.1126/science.167.3919.900

Terranova, J. I., Ferris, C. F., & Albers, H. E. (2017). Sex Differences in the Regulation of Offensive Aggression and Dominance by Arginine-Vasopressin. Front Endocrinol (Lausanne), 8, 308. 10.3389/fendo.2017.00308

Thompson, R. R., George, K., Walton, J. C., Orr, S. P., & Benson, J. (2006). Sex-specific influences of vasopressin on human social communication. Proc Natl Acad Sci U S A, 103(20), 7889–7894. 10.1073/pnas.0600406103

Tieger, T. (1980). On the biological basis of sex differences in aggression. Child Development, 51(4), 943–963. https://www.ncbi.nlm.nih.gov/pubmed/7471930

Tulogdi, A., Biro, L., Barsvari, B., Stankovic, M., Haller, J., & Toth, M. (2015). Neural mechanisms of predatory aggression in rats-implications for abnormal intraspecific aggression. Behav Brain Res, 283, 108–115. 10.1016/j.bbr.2015.01.030

Valenstein, E. S., Cox, V. C., & Kakolewski, J. W. (1970). Reexamination of the role of the hypothalamus in motivation. Psychol Rev, 77(1), 16–31. 10.1037/h0028581

Vertes, R. P., Linley, S. B., & Rojas, A. K. P. (2022). Structural and functional organization of the midline and intralaminar nuclei of the thalamus. Front Behav Neurosci, 16, 964644. 10.3389/fnbeh.2022.964644

Werhahn, J. E., Smigielski, L., Sacu, S., Mohl, S., Willinger, D., Naaijen, J., Mulder, L. M., Glennon, J. C., Hoekstra, P. J., Dietrich, A., Deters, R. K., Aggensteiner, P. M., Holz, N. E., Baumeister, S., Banaschewski, T., Saam, M. C., Schulze, U. M. E., Lythgoe, D. J., Sethi, A., … Brandeis, D. (2023). Different whole-brain functional connectivity correlates of reactive-proactive aggression and callous-unemotional traits in children and adolescents with disruptive behaviors. Neuroimage-Clinical, 40. ARTN 103542 10.1016/j.nicl.2023.103542

Williams, G., Bing, C., Cai, X. J., Harrold, J. A., King, P. J., & Liu, X. H. (2001). The hypothalamus and the control of energy homeostasis: different circuits, different purposes. Physiol Behav, 74(4-5), 683–701. 10.1016/s0031-9384(01)00612-6

Woodward, N. D., & Cascio, C. J. (2015). Resting-State Functional Connectivity in Psychiatric Disorders. JAMA Psychiatry, 72(8), 743–744. 10.1001/jamapsychiatry.2015.0484

Yang, T., Yang, C. F., Chizari, M. D., Maheswaranathan, N., Burke, K. J., Jr., Borius, M., Inoue, S., Chiang, M. C., Bender, K. J., Ganguli, S., & Shah, N. M. (2024). Social Control of Hypothalamus-Mediated Male Aggression. Neuron, 112(11), 1892. 10.1016/j.neuron.2024.05.010

Zhang, S., Hu, S., Chao, H. H., & Li, C. S. R. (2017). Hemispheric lateralization of resting-state functional connectivity of the ventral striatum: an exploratory study. Brain Structure & Function, 222(6), 2573–2583. 10.1007/s00429-016-1358-y

Zhang, S., & Li, C. R. (2018). Ventral striatal dysfunction in cocaine dependence – difference mapping for subregional resting state functional connectivity. Translational psychiatry, 8(1), 119. 10.1038/s41398-018-0164-0

Zhang, S., Wang, W., Zhornitsky, S., & Li, C. R. (2018). Resting State Functional Connectivity of the Lateral and Medial Hypothalamus in Cocaine Dependence: An Exploratory Study. Front Psychiatry, 9, 344. 10.3389/fpsyt.2018.00344

